# Telomerecat: A ploidy-agnostic method for estimating telomere length from whole genome sequencing data

**DOI:** 10.1101/139972

**Authors:** James HR Farmery, Mike L Smith, NIHR BioResource - Rare Diseases, Andy G Lynch

**Affiliations:** Cancer Research UK Cambridge Institute, University of Cambridge, Li Ka Shing Centre, Robinson Way, Cambridge CB2 0RE, UK; Genome Biology Unit, European Molecular Biology Laboratory (EMBL), 69117 Heidelberg, Germany.; NIHR BioResource - Rare Diseases, Cambridge University Hospitals, Cambridge Biomedical Campus, Cambridge, CB2 0QQ, UK

## Abstract

Telomere length is a risk factor in disease and the dynamics of telomere length are crucial to our understanding of cell replication and vitality. The proliferation of whole genome sequencing represents an unprecedented opportunity to glean new insights into telomere biology on a previously unimaginable scale. To this end, a number of approaches for estimating telomere length from whole-genome sequencing data have been proposed. Here we present Telomerecat, a novel approach to the estimation of telomere length. Previous methods have been dependent on the number of telomeres present in a cell being known, which may be problematic when analysing aneuploid cancer data and non-human samples. Telomerecat is designed to be agnostic to the number of telomeres present, making it suited for the purpose of estimating telomere length in cancer studies. Telomerecat also accounts for interstitial telomeric reads and presents a novel approach to dealing with sequencing errors. We show that Telomerecat performs well at telomere length estimation when compared to leading experimental and computational methods. Furthermore, we show that it detects expected patterns in longitudinal data, technical replicates, and cross-species comparisons. We also apply the method to a cancer cell data, uncovering an interesting relationship with the underlying telomerase genotype.

## Introduction

Telomeres are the ribonucleoprotein structures at the ends of chromosomes. They are multifunctional regions of the genome that serve to protect coding DNA from the shortening process inherent in cell replication; that act as a molecular clock; and that shield the ends of chromosomes from the DNA damage response^1^. In humans, the telomere is an extremely repetitive region of the genome comprised of the nucleotide hexamer: (*TTAGGG*)_*n*_. Telomere length is both a driving force in tumour aetiology and a risk factor for cancer and other diseases^2,3^.

In this study we present Telomerecat, the first tool designed specifically to estimate mean telomere length from cancer whole genome sequencing (WGS) data. There have been previous approaches to using WGS data to say something about telomeres. Castle et *al.* provided a proof of concept in 20104, and this was refined by the first group to use such an approach in earnest^5^. Ding *et al*.^6^ published the first fully-fledged method for estimating length rather than just telomere content, with the accompanying tool ‘TelSeq’. Their study was also the first time a computational method had been validated against an established experimental method.

TelSeq assumes a fixed number of chromosomes when estimating telomere length and so makes no allowance for aneuploidy. Nevertheless, as the strongest available tool there are several examples of TelSeq being used to analyse cancer datasets^7,8^ Ṅotably a recent pan-cancer analysis made use of the TelSeq tool^9^. While generally sound, such analyses are vulnerable to misinterpretation in the event of systematic differences in aneuploidy (as may be the case when comparing different cancer types). Indeed, recurrent somatic copy number alterations involving the telomere were observed in all cancer types studied in a pan-cancer study of Cancer Genome Atlas data^10^.

Where such changes (suggestive of aneuploidy) occur, cells will likely be left with an altered number of telomeres. Accordingly the quantity (and proportion) of telomere sequence within the sample is altered, even if the mean length of telomeres is unaltered. Thus if we observe more telomere sequence in a cancer sample, we do not know if this is due to longer telomeres.

Two other tools of note have been published: TelomereHunter and Computel. TelomereHunter^11^ reports telomere content rather than telomere length, and so does not provide a direct comparison. TelomereHunter classifies reads based on their mapping location within the parent BAM file and outputs statistics relating to variations of the canonic telomere hexamer. Computel^12^ does allow the user to specify the number of telomeres present, but since this is unknown (and cannot safely be inferred from copy-number profiles or ploidy statistics) it again does not provide a direct comparison. Since TelSeq is more frequently used in the literature, has greater experimental validation than Computel, and a recent comparison study^13^ did not find that the greater convenience of TelSeq was at the cost of poorer performance, we take TelSeq as the representative of current methods in our comparisons.

Telomerecat accounts for aneuploidy as an inherent part of the method, without relying on knowledge of the number of telomeres present, and so avoids such potential misinterpretation of results. Another source of error for tools of this nature can arise from stretches of the TTAGGG repeat sequence that appear in the human genome distal from the actual telomeres: so-called Interstitial telomeric repeats (‘ITRs’)^14^. As well as a consideration of aneuploidy, Telomerecat estimates and corrects for the number of ITR-originating reads without consideration for how these highly repetitive reads are aligned to a reference genome. This removes any reliance on upstream preprocessing by a sequence aligner for the removal of ITR reads and increases the wide applicability of the method.

A third potential hindrance is that it is difficult to define the end of the telomere precisely based solely on genomic sequence (explicit information about DNA secondary structures and the locations of bound proteins having been lost). The subtelomere is composed of subtelomeric repeat sequences and segmental duplicates, interspersed by canonic telomere repeats^15^. These subtelomeric repeat sequences can look much like the telomere but with the addition of sequencing errors. Too strict a definition of telomere as being the region of TTAGGG repeats would be hostage to genuine variations, sequencing errors, and somatic mutations.

Telomere length is therefore necessarily a subjective measure, consistent only within the method used. Accordingly there may be an off-set in comparisons with other methods. Even ‘gold standard’ laboratory methods for measuring telomere lengths may have their own biases in this regard^16^.

Moreover, differences in patterns of sequencing error have the potential to lead to inconsistency between samples even if using the same method. To this end, Telomerecat includes a novel method for correcting sequencing error in telomere sequencing reads. This model automatically adapts to differing error across sequencing preparations.

Telomerecat is an open source tool, the code is available from https://github.com/jhrf/telomerecat. Full installation and usage documentation is available at https://telomerecat.readthedocs.io

## Results

### Validation in presumed-diploid blood samples

To verify that Telomerecat is able to identify telomere length within WGS samples, we compared the algorithm to an established experimental method (mean terminal restriction fragment Southern blot experiment (mTRF)) and the leading computational method (TelSeq). Blood samples were taken from 260 adult females as part of the TwinsUK10K study, WGS and mTRF were conducted on each sample (described previously^17 18^). The donor’s age at sample collection is also recorded for each sample. Since absolute agreement is not expected, we consider correlations between the methods. The results of the comparisons are shown in Table 1 and in Figure 1.

**Table 1.** Results for the comparisons between Telomerecat, TelSeq, mTRF and Donor Age. Pearson correlation was used for each comparison.

|  | Telomerecat | TelSeq | mTRF |
| --- | --- | --- | --- |
| TelSeq | $\rho = 0.631$ | - | - |
| mTRF | $\rho = 0.618$ | $\rho = 0.583$ | - |
| Donor Age | $\rho = -0.306$ | $\rho = -0.239$ | $\rho = -0.321$ |

**Figure 1.**
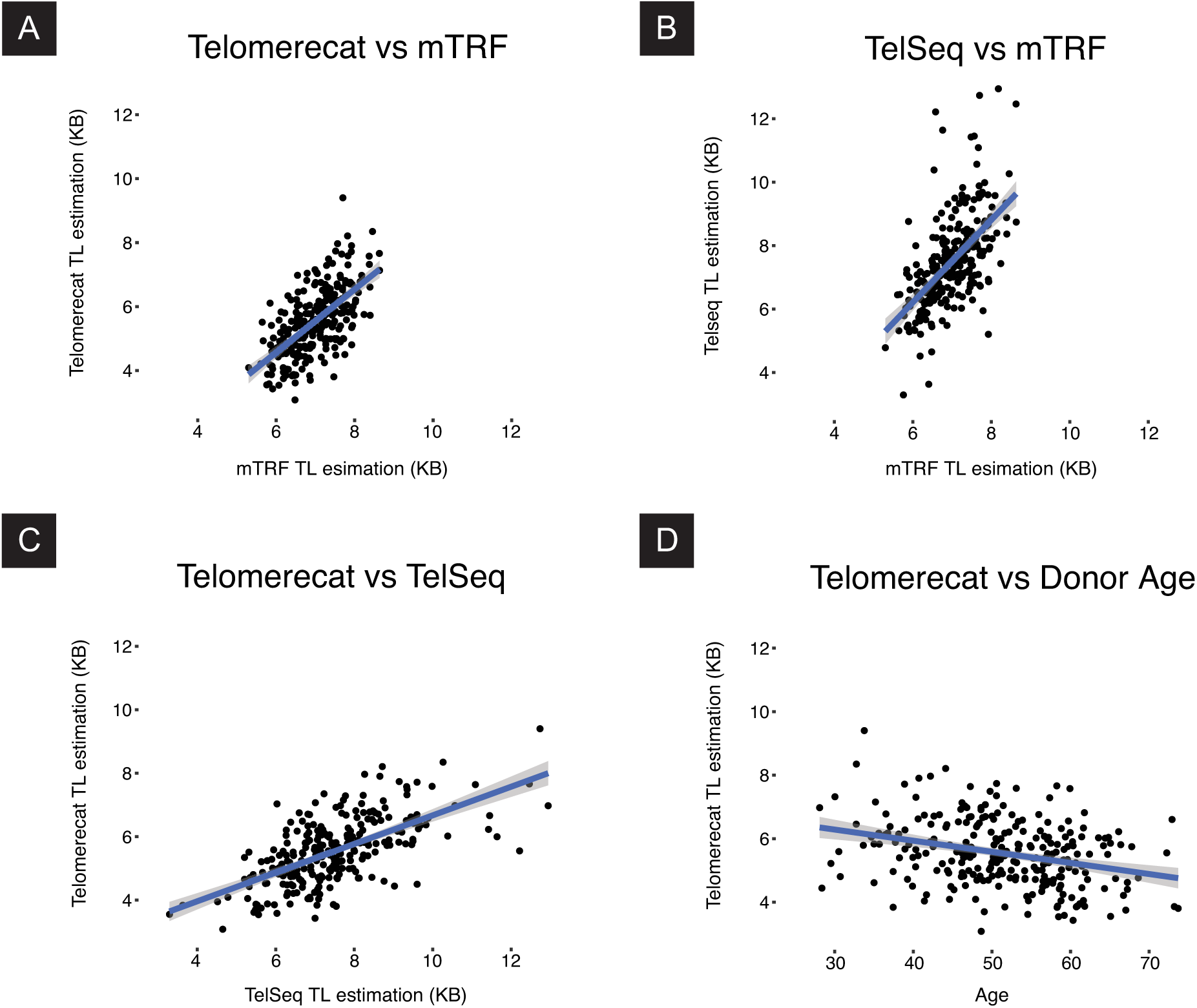
Scatter plots describing the relationship between Telomerecat, mTRF, and TelSeq estimates of telomere length (TL).

We observe that the best correlation is between the the two computational methods at *ρ* = .631. The next best correlation was between mTRF and Telomerecat indicating that Telomerecat agrees with the established experimental method. Both Telomerecat and TelSeq correlate well with mTRF indicating that both tools are providing realistic estimates of telomere length. The extent that Telomerecat correlates with mTRF is in line with correlations previously observed between other experimental methods and mTRF^16^.

Telomerecat estimates telomere length that is shorter, on average, than TelSeq. At least part of this disparity may be due to Telomerecat’s active filtering of reads from ITRs. Telomerecat finds that, on average 7% of telomeric read-pairs identified are from ITRs.

Telomerecat was able to identify a correlation with age only slightly weaker than that of mTRF, a strong indicator that we are capturing genuine information about telomere lengths.

### Application to a longitudinal MSC data set

We applied Telomerecat to a set of WGS samples from a mesenchymal stem cell (MSC) experiment described previously^19^. Mesenchymal stem cells are multipotent stromal cells commonly located in bone marrow^20^. The experiment constituted six WGS samples: an in vivo MSC sample from a healthy 31 year old male, three passaged MSC samples (P1,P8 and P13) and two induced pluripotent stem cell (iPSC) samples.

MSCs are unusual amongst mature human stem cells as they do not express any measurable amount of telomerase^21^. Accordingly, telomere length attrition has been described in MSC passage experiments^22^. Conversely, iPSC cells have been shown to exhibit heightened telomerase expression^23^. We hypothesised that telomere length would shorten across the passaged MSC samples and lengthen within the iPSC samples.

We applied Telomerecat and TelSeq to the aforementioned MSC WGS data. The results are shown in Figure 2. Telomerecat identifies telomere shortening across the passaged samples, as expected. Telomerecat estimates that between P1 and P13 the average telomere length was shortened by 2.5KB, at a rate of approximately 0.2KB per passage. Furthermore, we see that Telomerecat identifies long telomere length in the the two iPSC samples. We also note that TelSeq fails to identify the expected telomere dynamics.

**Figure 2.**
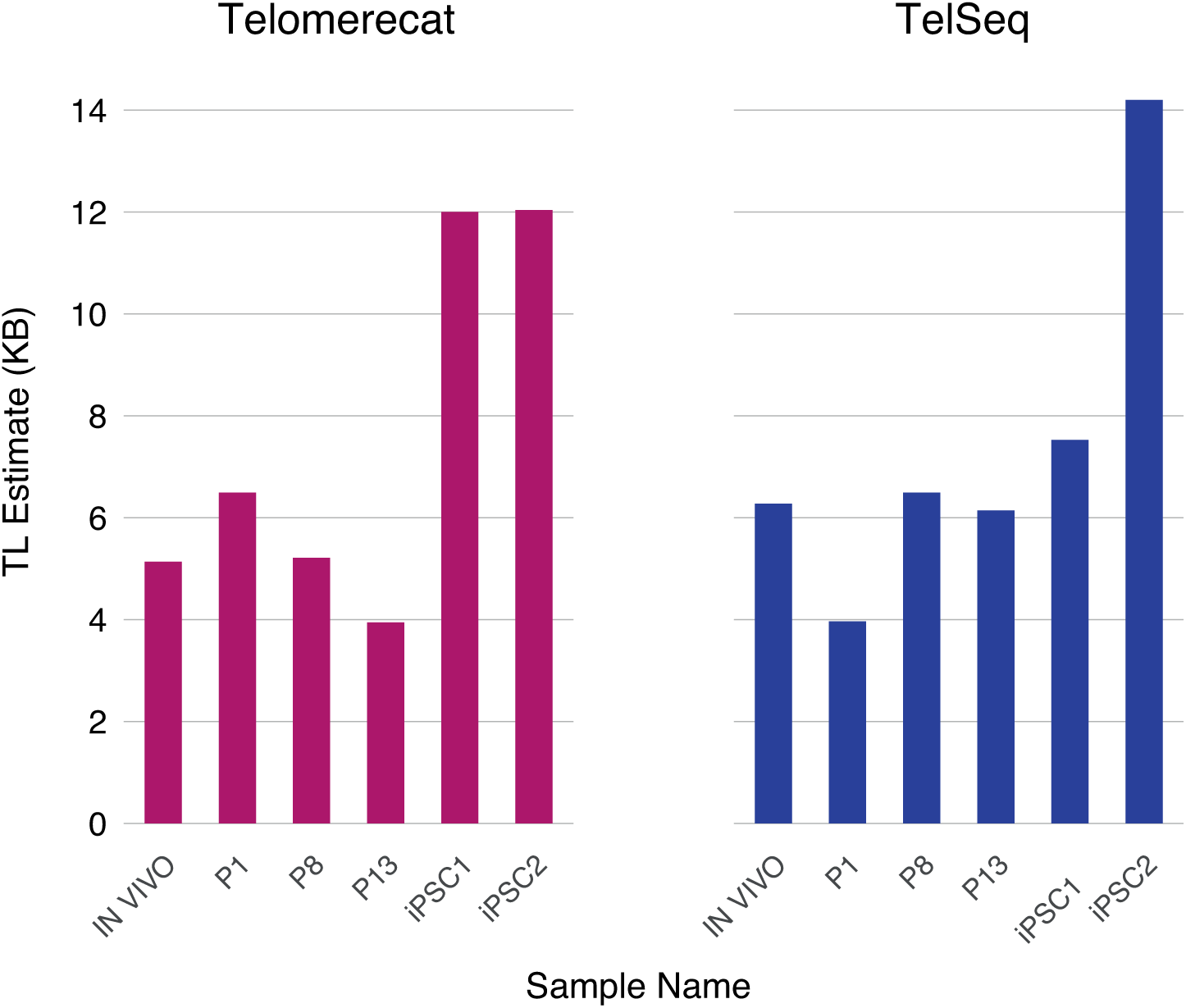
This figure shows estimates for the MSC samples produced by Telomerecat (left) and TelSeq (right). We expect to see a decrease in telomere length with additional passaging (P1 to P13), but consistent high telomere lengths in the two iPSC samples (iPSC1 and iPSC2)

### Application to a cancer dataset

After establishing that Telomerecat performs well in diploid samples, we demonstrated that it can also be applied to cancer samples. We applied Telomerecat to a data set comprised of samples from four donors suffering from Hepatocellular carcinoma (HCC)^24^. Primary HCC cells were extracted from each donor in that study. These primary cells were cultured to create cell lines. Samples of the primary cells in vitro, an early passage and a late passage were taken for sequencing. Table 2 lists the exact passage number for each sample.

**Table 2.** Patients in the HCC study

|  | CLC11 | CLC13 | CLC16 | CLC5 |
| --- | --- | --- | --- | --- |
| Early Passage Count | 6 | 3 | 3 | 7 |
| Late Passage Count | 24 | 18 | 21 | 27 |
| TERT Promoter Mutation | No | Yes | Yes | Yes |
| TERT Amplification | Yes | Yes | No | No |
| HBV Integration | Yes | No | No | No |

Figure 3 shows the results of applying Telomerecat to the HCC cohort. We observe two telomere length phenotypes across the four donors. CLC11 and CLC13 show a telomere length that is not altered across the passage process. By contrast, in CLC16 and CLC5 we see that telomere length increases across the passaged samples. Z. Qiu *et. al* report that all four samples contain corruptions in the TERT gene as shown in Table 2. It is interesting to note that CLC16 and CLC5 share both a TERT genotype and telomere length phenotype.

**Figure 3.**
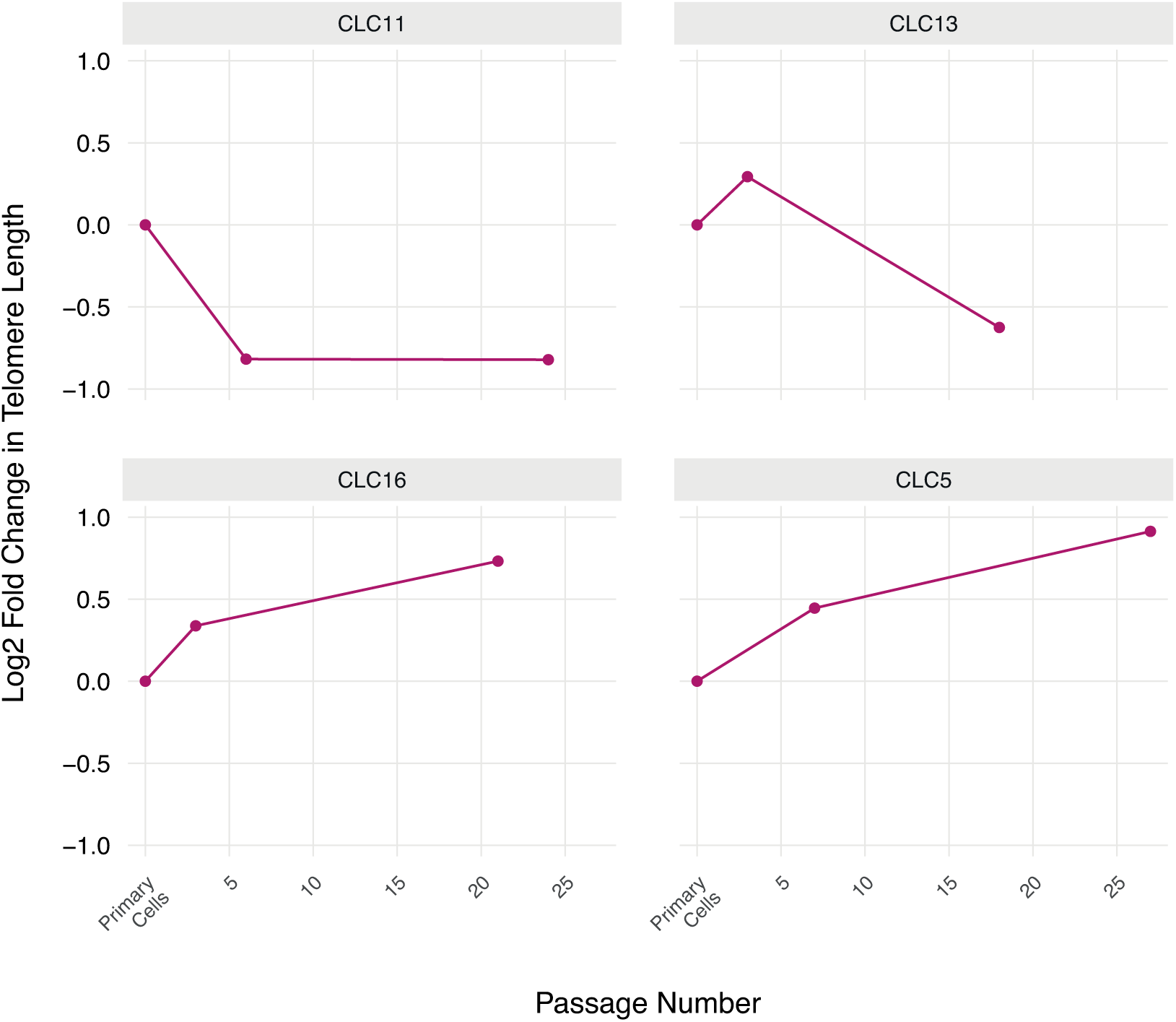
Telomerecat estimates for the HCC cell line dataset. Results are shown as log2 fold change in relation to the Primary Cell telomere length

Previous studies suggest that the presence of TERT promoter mutations and HBV Integration increases TERT expression^25,26^. However it is not clear that heightened expression is indicative of longer telomere lengths. Indeed, HCC tumours generally have shorter telomeres than adjacent normal cells^27^.

### Application to a set of technical replicates

We have also tested Telomerecat on pairs of WGS technical replicates from the NIHR BioResource - Rare Diseases study. Telomerecat was applied to 93 samples of DNA extracted from whole blood. For each participant two samples were taken. Each sample was sequenced on either the HiSeq2000 or HiSeqX platform. We observe cases in this cohort where samples from the same participant were sequenced on the same technology and where samples were sequenced on different technologies.

A sound approach to telomere-length estimation will be reproducible across duplicate samples. After accounting for batch effects relating to choice of platform, Telomerecat achieves good agreement between duplicate pairs, as shown in Figure 4.

**Figure 4.**
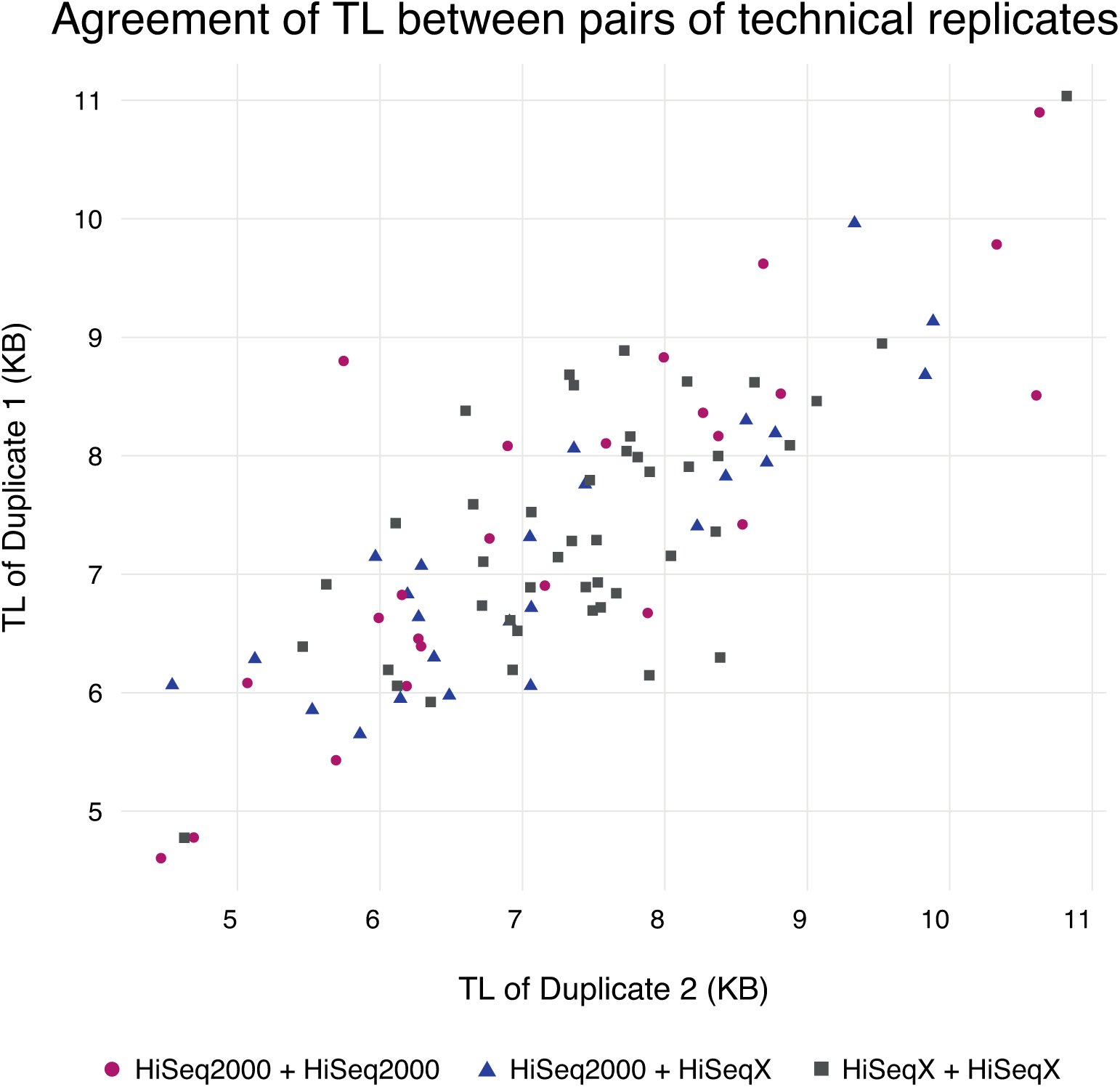
A plot of telomere length (TL) estimates for technical duplicate pairs. Colours correspond to the sequencing platform of each sample in the pair.

### Application to mouse samples

Mouse telomeres are known to be longer than human telomeres^28^. However, telomere length is known to vary across different mouse strains. We applied Telomerecat to 10 samples from the Mouse Genomes Project^29^. Telomerecat identifies a range of telomere lengths, most of which are substantially greater than estimates from human samples. The estimates for the mouse samples, as well as two human samples for comparison, are shown in Figure 5. TelSeq was not applied as the tool is specifically tailored to the human genome.

**Figure 5.**
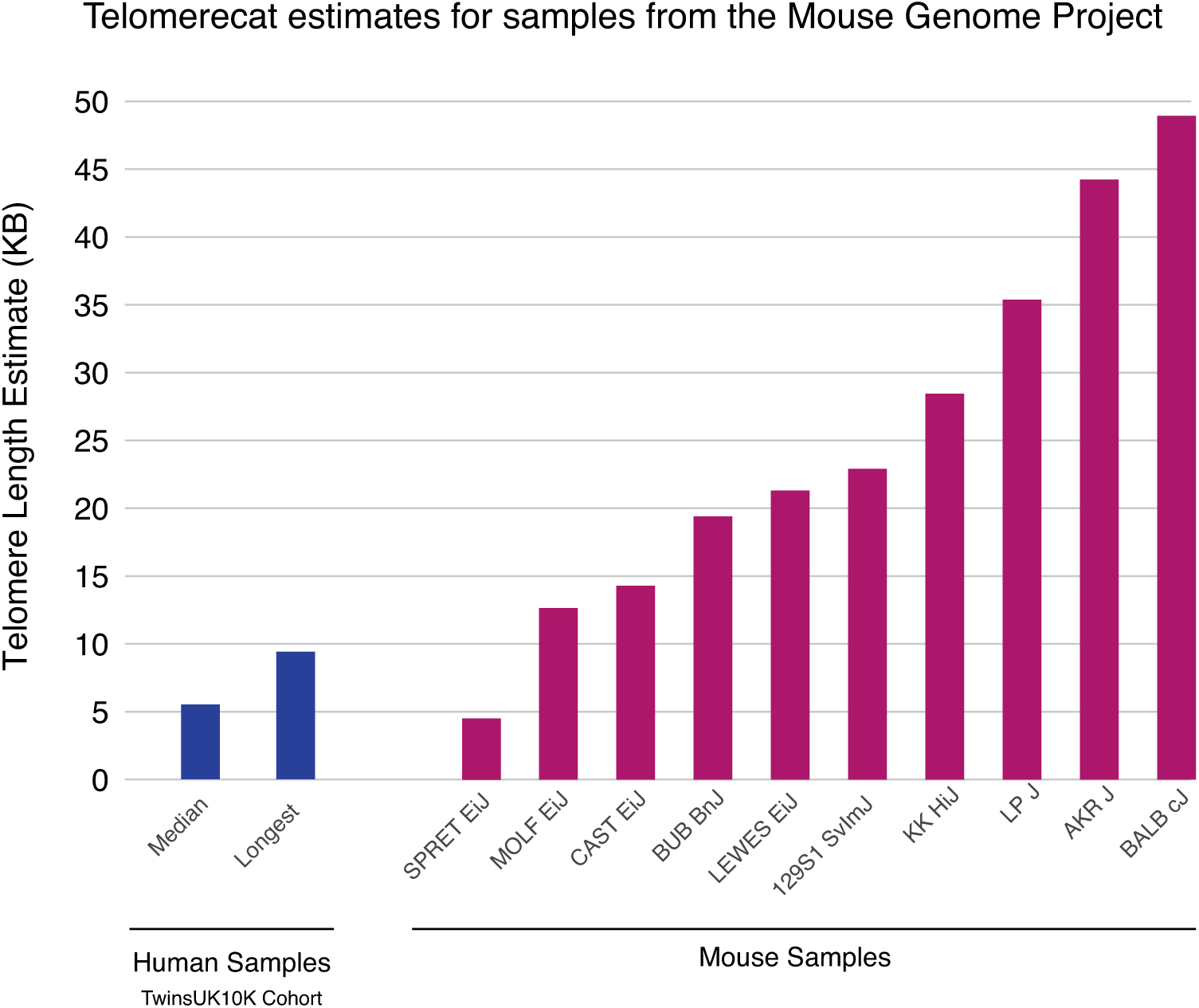
Telomere length estimates by Telomerecat for 10 mouse samples from the Mouse Genomes Project

Telomerecat identifies a range of telomere lengths for the mice, almost all of the lengths are substantially longer than the longest human telomeres in the TwinsUK10K cohort. Additionally, we note that two of the samples with the shortest estimates - CAST Eij and SPRET Eij - have been identified as having comparatively short telomeres^30–32^. We also note that previous studies have identified the BALB cJ mouse strain as having long telomeres^32^.

### A comparison of running time and resource allocation

Benchmarking was conducted on a MacPro desktop computer with 2x 2.93 Ghz Quadcore Intel Xeon processors and 16GB of 1066Mhz DDR3 memory. The results of benchmarking for the Telomerecat and TelSeq tools can be found in Table 3. Benchmarking was conducted on QTL190044 from the TwinsUK10K cohort. The results displayed are the average from the three runs.

**Table 3.** Benchmarking results for Telomerecat and TelSeq

|  | Telomerecat | TelSeq |
| --- | --- | --- |
| Time Taken (seconds) | 756 | 3894 |
| Reads per hour | $1.562 \times 10^9$ | $3.299 \times 10^8$ |
| Max. Processor Usage (%) | 537.6 | 96.8 |
| Avg. Processor Usage (%) | 356.8 | 80 |
| Max. Memory Usage (GB) | 1.9 | 0.104 |
| Avg. Memory Usage (GB) | 1.3 | 0.037 |

## Discussion

Here we have demonstrated and validated a novel approach to estimating telomere length from WGS data. Importantly, Telomerecat is the first tool designed to be applicable to cancer experiments as it does not assume a given number of telomeres.

Core to Telomerecat’s estimation process is the ratio between read-pairs that lie within the telomere and read-pairs that span the telomere boundary. Observing reads on the boundary between telomere and subtelomere provides a quantification of telomere numbers through which we normalize the telomere lengths. Where other samples always assume that more telomere reads mean longer telomere, Telomerecat is able to account for the fact that there may actually be more individual telomeres.

We have validated Telomerecat by showing that it correlates with existing computational and experimental methods as well as with sample donor age. mTRF itself provides an imperfect measure of telomere length and, from correlations with age, it seems that computational methods may be capturing as much information as that approach.

WGS-based methods will naturally be more accurate as the depth of sequencing increases.

Much of the inaccuracy in the estimates of the TwinsUK10K data may be attributable to the relatively low coverage of the WGS data. At low coverage, Telomerecat’s estimate of the number of reads crossing the boundary is less certain. As coverage at the boundary decreases and the observed read counts for each individual sample become less certain Telomerecat relies more on the cohort error adjustment (discussed in the methods section). With higher coverage we would expect even better agreement between Telomerecat and the other methods for diploid cells.

By applying Telomerecat to the duplicate blood samples we have demonstrated Telomerecat’s ability to generate meaningful results on two of the most popular Illumina paired-end platforms. As well as confirming the reliability of Telomerecat’s telomere length estimates, this shows that the estimates are robust to sequencing batches once batch effects are accounted for.

Amongst the most striking results presented here is the estimation of telomere length across MSC cell line passaged data. Telomerecat identifies a clear deterioration of telomere length across the passaged cells and an increase of telomere length in the iPSC samples, in which telomerase had been reactivated. Notably, TelSeq fails to identify this pattern.

On observing TelSeq’s output, we see that the most likely reason for its failure to observe the expected telomere dynamics is in the GC correction part of the algorithm (see Supplementary Information). This indicates that the relationship between coverage at locations where genomic GC is identical to telomere and actual telomere, on which TelSeq relies, may not always be consistent across experiments.

We have presented the first application of a WGS telomere length estimation approach to data derived from non human samples; Telomerecat’s agnosticism to telomere numbers provides a natural advantage here also. As expected, Telomerecat identifies long telomere length in most of the mice samples. Pleasingly, Telomerecat is concordant with the literature in demonstrating the short telomeres in CAST EiJ and SPRET cJ samples and long telomeres in BALB cJ.

Telomerecat tends to report shorter telomere length than other methods, both computational and experimental. There will be several contributing factors, including disagreement over the definition of the telomere/sub-telomere boundary, and the stringency for categorizing read-pairs as being telomeric. One clear contributing factor in the comparison of computational methods will be Telomerecat’s exclusion of ITR read-pairs, typically contributing 4% to 10% of apparently telomeric read-pairs.

We have also demonstrated that Telomerecat can be run quickly (five times faster than TelSeq for our example). Telomerecat is able to process samples quickly as it is built on a parallel BAM processing framework - parabam^33^ - and thus uses multiple processing cores at all stages of the analysis. Telomerecat promotes reproducible research by generating subsets of reads from which telomere length estimates can be generated. We hope that these smaller file will be more easily stored and transferred allowing researchers to regenerate estimates without the need to process the cumbersome original BAM files.

Finally, we have demonstrated the application to a cancer WGS dataset: Telomerecat’s raison d’être. We see that Telomerecat identifies differing telomere phenotypes across four passage experiments. Intriguingly the two experiments with the most similar telomere length phenotype have an identical underlying TERT corruption.

## Methods

### Overview

Telomerecat functions as three discrete operations: TELBAM generation, read categorisation and length estimation. A flowchart depicting the method is given in Figure 6.

**Figure 6.**
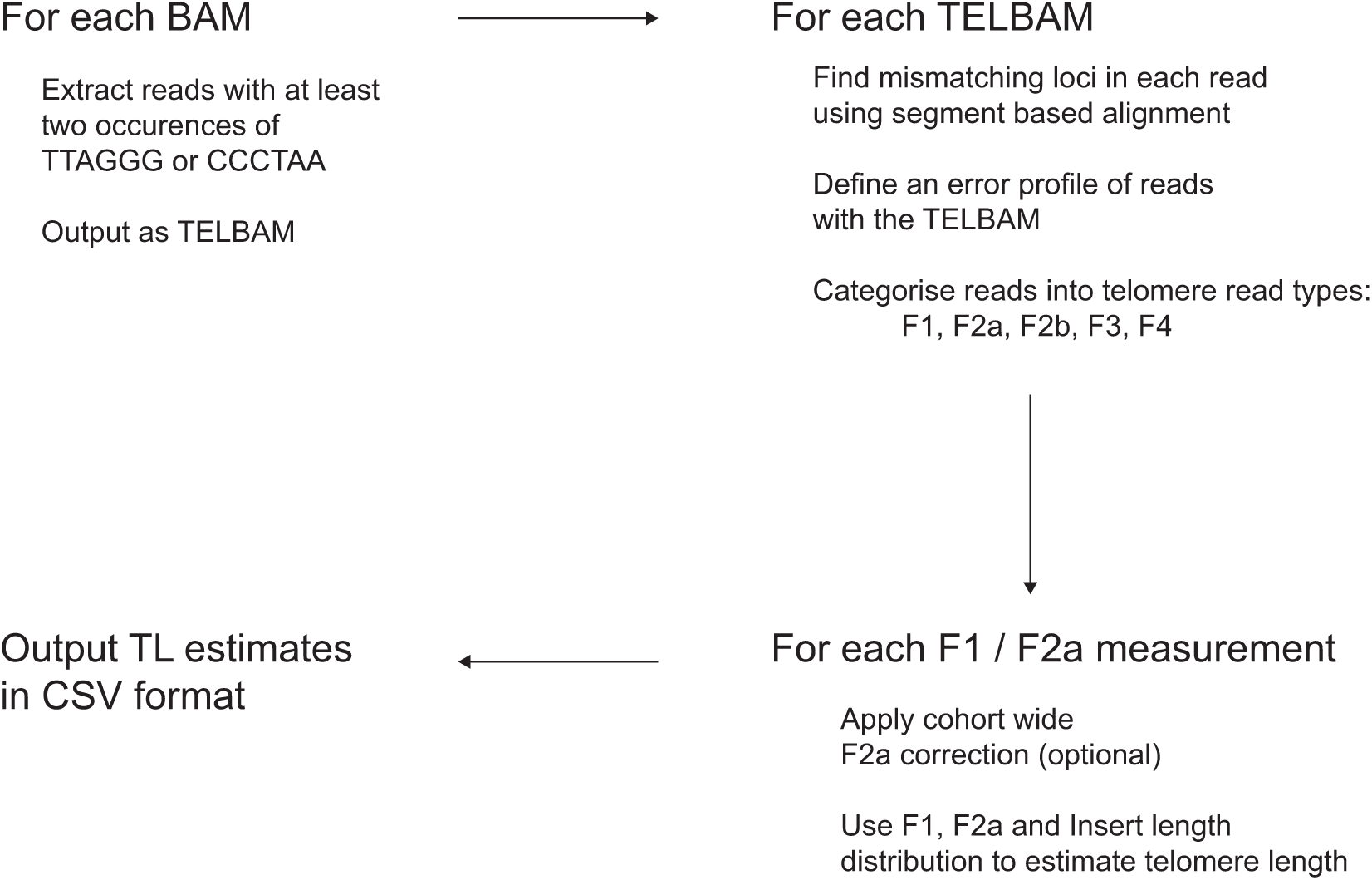
An overview of the Telomerecat length estimation process

First, we collect a relevant subset of reads and their pairs from a BAM file. This subset is referred to as a TELBAM and consists of read pairs where one end of a read pair has two occurrences of the telomeric hexamer. This read subsetting operation is expedited by using the parallel processing framework parabam^33^. We observe that TELBAMs contain less than one ten-thousandth of the reads from an input BAM file.

Next we categorise read pairs according to their sequence composition and orientation on the genome. The telomere length estimate is informed by a ratio of complete telomere read pairs to read pairs on the boundary between telomere and subtelomere. In order to differentiate between the various type of telomere read we must first understand how reads differ from the telomere sequence and whether these differences are genuine biological perturbations or the result of sequencing error.

Lastly, we use the ratio of complete to boundary read-pairs in conjunction with insert length distribution to estimate the underlying telomere length that produced the observed complete to boundary ratio.

### Defining error in telomere reads

Key to the process of identifying sequencing error is identifying loci within reads that do not match the expected telomere sequence. We shall refer to these as “mismatching loci”. Telomeres are extremely repetitive stretches of DNA. This repetition of sequence allows us to imagine a hypothetical telomere sequence and then to compare reads to the hypothetical sequence to find differences. In order to account for insertions and deletions in the sequencing reads (both biological and as a result of sequencing error) we use a method of fragmentary local alignment. Reads that suffer few mismatches, and those mismatches at loci with low Phred scores, likely represent complete telomere sequences.

Since mismatch loci that represent sequencing errors should be associated with lower Phred scores, we first observe the empirical joint distribution of Phred scores at mismatching loci, and number of mismatching loci across the BAM file (Figure 8A) before constructing the equivalent distribution for loci chosen at random within the reads (Figure 8B). We find that reads with few mismatches and low Phred scores (complete telomere sequences suffering from sequencing error) are over-represented in the empirical data set.

We define *P_max_* and *P_min_* as the global maximum and minimum observed Phred score across all reads, and (*L*) as the read length used.

We let *N* represent the total number of reads in the TELBAM such that {0, 1, *n*,…,*N* − 1} are indices representing each read. Values associated with the *n^th^* read are denoted with a superscript (*n*). For example, the vector of Phred scores associated with the L locations in read n is denoted 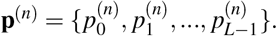. For the *n^th^* read, let *m*^(*n*)^ be a random vector in the space {0, 1}^*L*^ such that a 1 is found at each loci in the read that does not agree with the telomere sequence. In the case that the sequence is comprised of perfect telomere sequence then the vector should sum to zero. The method for obtaining *m*^(*n*)^ via an fragmentary alignment method is shown in Figure 7.

**Figure 7.**
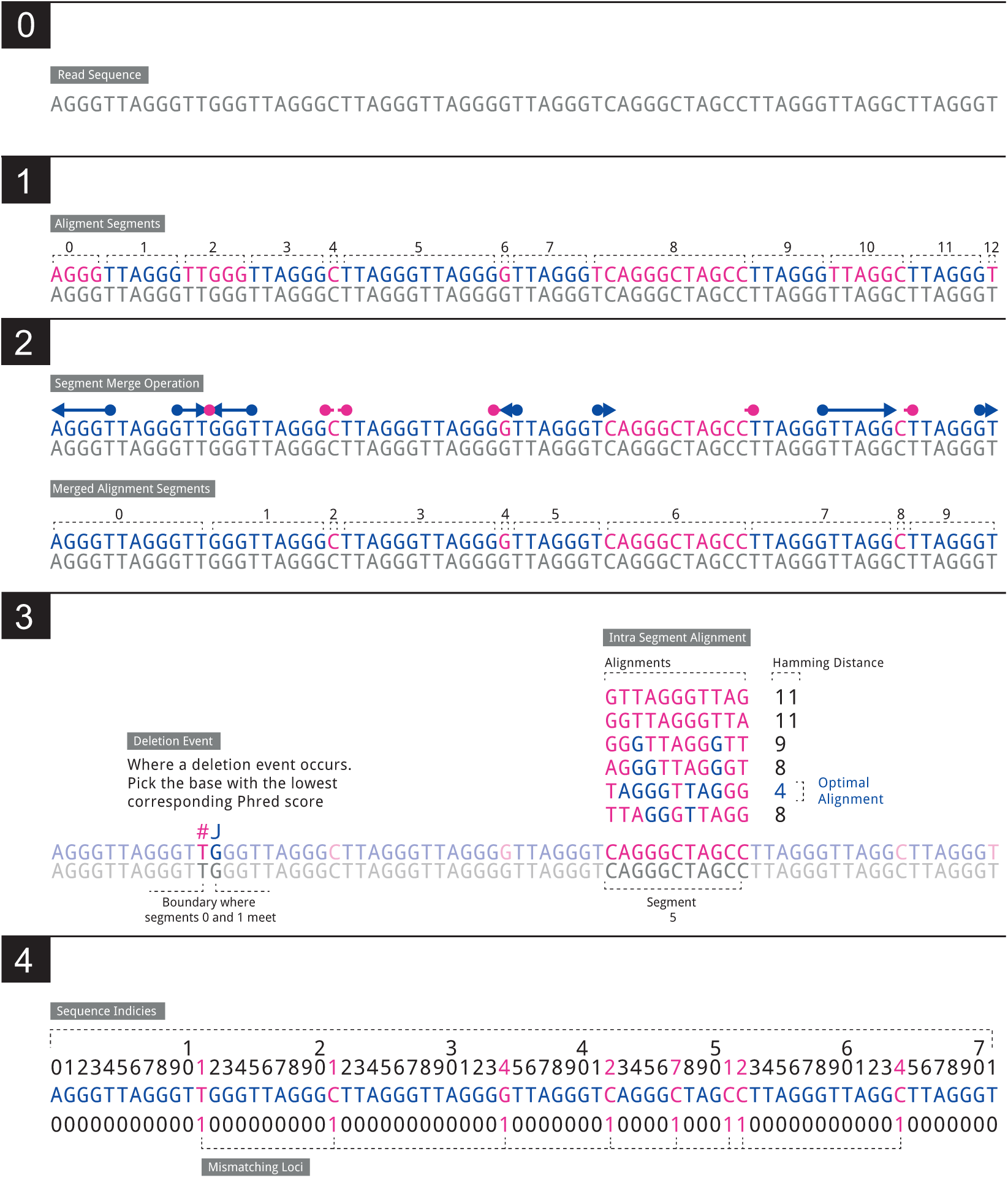
The algorithm that determines the indices of divergence from the telomere sequence. **0**: We observe a sequencing read **1**: We split the read into ‘segments’ (11 in total in our example) such that each segment is a substring of the original sequence and that every other segment consists of unbroken telomere sequence. In our example we see that segments 1,3,5,7,9,11 contain unbroken telomere sequence. **2**: Each segment containing a telomere hexamer is ‘expanded’ to capture the full extent of the surrounding telomere sequence. The number of segments is reduced by 2. **3**: When two segments both containing the telomere hexamer are adjacent after Step 2 this indicates a deletion event. We take the loci with the lowest corresponding Phred score. For any segment that does not contain a telomere hexamer and where the length of the segment is greater or equal to 4 apply we conduct a basic alignment of all possible telomere offset telomere sequences. The telomere sequence with the lowest Hamming distance is taken as a local alignment for that segment. Where two alignments are equal the one with the lowest average Phred score is preferred. **4**: Sequence loci that are not in a complete hexamer or were mismatched in the Hamming alignment step are taken as mismatching loci. **m** for this example is given in the final line of the diagram.

Then define *z^n^* (the number of mismatches for read n), and *λ^n^* (the average Phred score at mismatches in read n)) as:

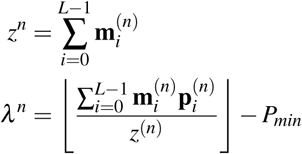

We then define an indicator function

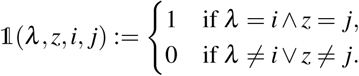

So that a matrix **X** takes the form,

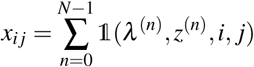

Where *i* ∈ {0,…,*P_max_* – *P_min_*} and *j* ∈ {0,…,*L*−1}. Thus each *x_ij_* in **X** records the number of reads with the relevant *λ* and *z* contained within the TELBAM and is depicted in Figure 8A.

**Figure 8.**
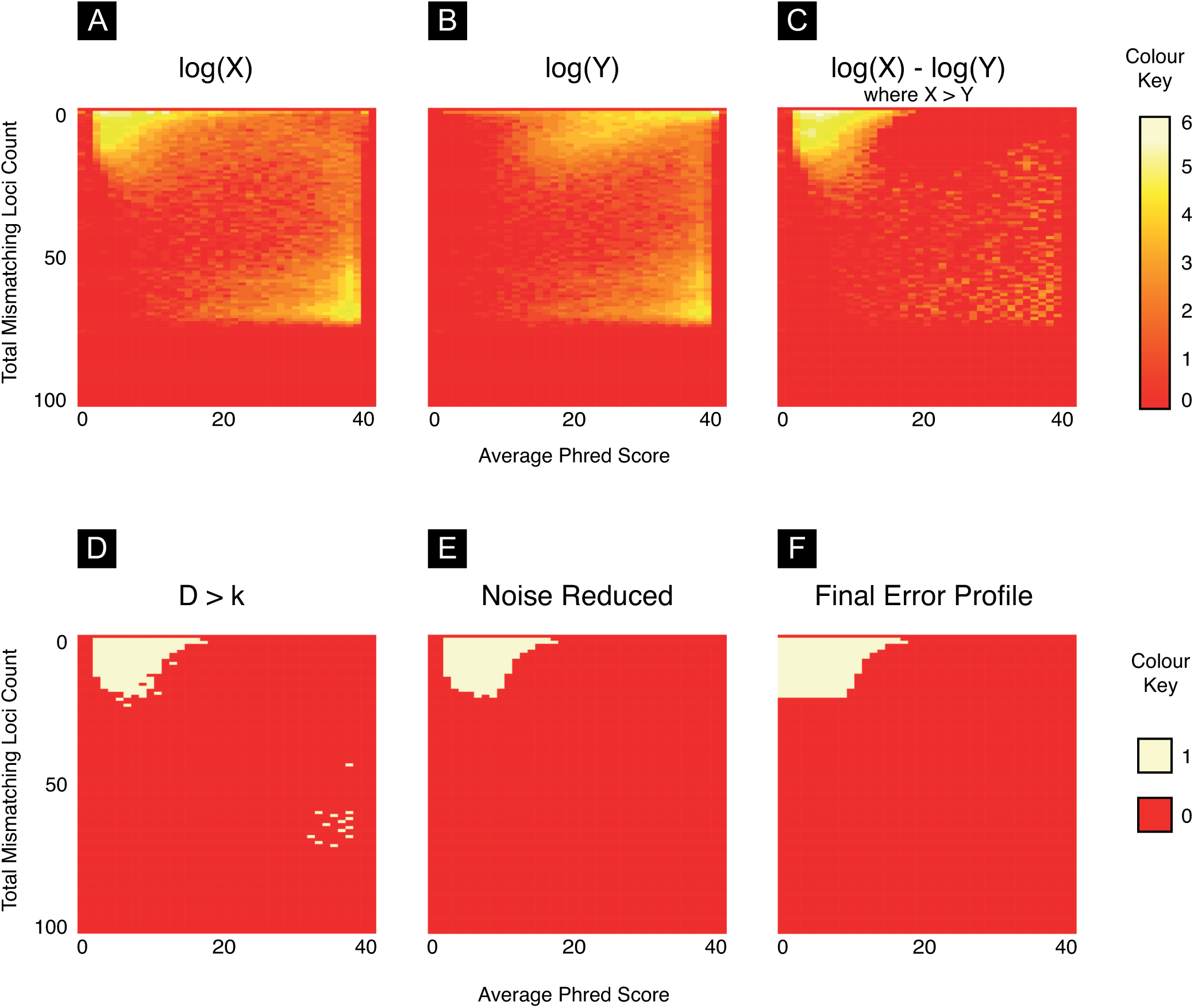
***A***: A heatmap of the joint distribution of Phred scores a mismatching loci and the number of mismatching loci (**X**). The intensities in the top left corner of the heatmap indicate an association between fewer mismatches and lower phred scores. We observe that the maximum mismatching loci is commonly ~75% of the read length. This effect is caused by non-telomere reads match a the telomere sequence simply by chance ***B***: A heatmap of the joint distribution of random loci in reads and the associated phred score (**Y**). We note that the joint distirubtion of reads in the upper half of the matrix is different to that in **X** while the lower portion is identical. ***C***: The difference between **X** and **Y**. Referred to as **D** in the text. ***D***: A binary heatmap showing all cells in **D** that are greater than the threshold *k*. We note the preponderance of cells in the upper left hand corner of the figure ***E***: We remove noise from the figure using the methods detailed in (Supplementary Algorithm 1) ***F***: We apply a final rule to ensure cells associated with low Phred scores are captured in the error profile (Supplementary Algorithm 2)

Where **X** captures information about the average Phred score (*λ*^(*n*)^) at *z*^(*n*)^ mismatching loci, we seek to create an equivalent matrix **Y** about the average Phred score at *z*(*n*) random loci in the *n^th^* read.

For the *n^th^* read, let *r*^(*n*)^ be a random vector in the space {0, 1}^*L*^ such that 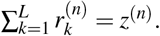. That is, a vector for which the non-zero entries identify *z*^(*n*)^ random loci within the read.

So that,

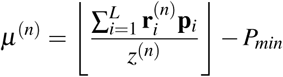

Thus,

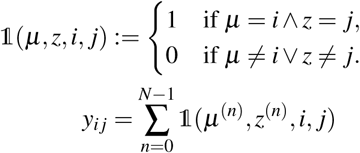

As before, *i* ∈ {0,…,*P_max_* − *P_min_*} and *j* ∈ {0,…,*L* − 1}.

When we plot the matrices **X** (Figure 8A) and **Y** (Figure 8B) as heat maps we typically see that there is a striking difference in their composition. The heatmap for **X** shows an intensity in the upper left hand corner pertaining to reads with low Phred scores at mismatching loci. This hotspot is missing from the **Y** heatmap. We interpret this region as representing telomere reads affected by sequencing error that we wish to capture in our length estimation process.

We find the difference between the two matrices:

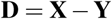

We plot values of **D** > 0 as a heatmap in 8C. To capture cells that contain more reads than we would expect at random we define a mask **E**. **E** is defined such that:

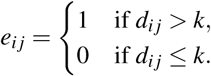

Where *k* is max{**D**_*ij*_} for all values where 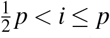 and 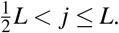. This matrix is depicted as a heatmap in Figure 8D.

We note that the mask depicted in Figure 8D has gaps that appear as a result of using k as a threshold. We apply the procedure detailed in Supplementary Algorithm 1 in order to remove noise from the error profile. The results of applying this procedure are shown in Figure 8E. We conclude by applying the operation described in Supplementary Algorithm 2 and shown in Figure 8F. This is the final matrix and is provided to the read classification procedure shown in Supplementary Algorithm 3 as **E**. All reads falling within the area by the error profile are counted as fully telomeric suffering from sequencing error.

Our definitive definition of a fully telomeric read is a read where 90% of the the sequence is telomere or the read falls into the error profile (See Supplementary Algorithm 3). In practice we observe that using a threshold above 90% leads to decreased accuracy. It is possible that this is indicative of genuine telomere heterogeneity but further study is required to understand this phenomenon.

### Categorising telomere read types

Once we have adequately described sequencing error we now classify each read-pair. In this section we describe the step that allows us to sort read-pairs into ‘complete’ read-pairs (denoted F1 reads in Figure 9 - both reads of the pair lying wholly within the telomere) and boundary (F2a - exactly one read of the pair lying wholly within the telomere) reads.

**Figure 9.**
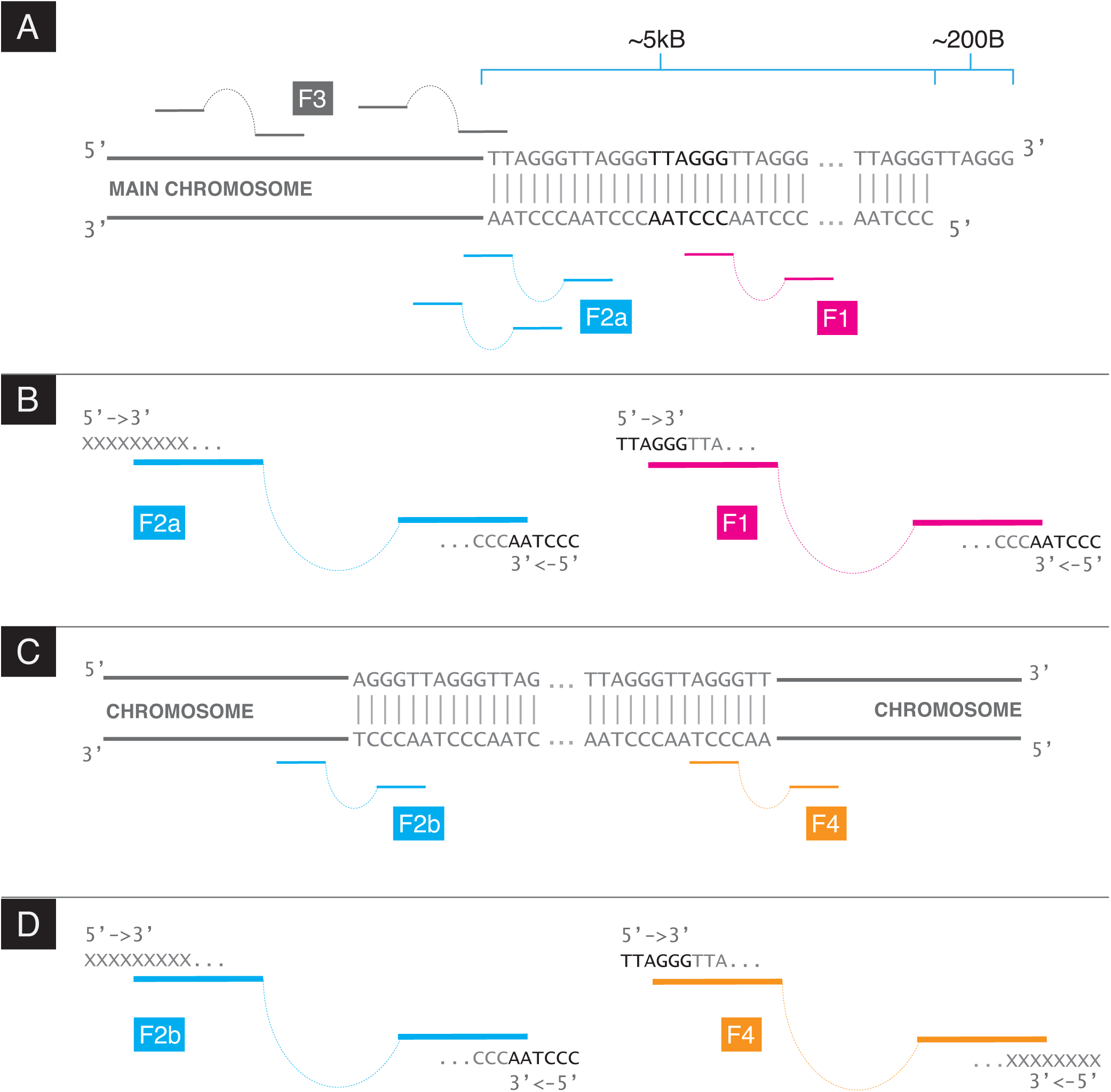
**A**: The read-pair types at the boundary between telomere and subtelomere. F2a reads stem from the boundary whereas F1 reads stem from anywhere within the telomere proper. F3 are reads where neither read in the pair is complete telomere **B**: Detail of the F1 and F2a read types. F1 read-pairs are comprised of two complete telomere reads. F2a read-pairs are comprised of a read-pair where one read is complete telomere and the other is not. Crucially, the complete telomere read is comprised of CCCTAA **C**: The read-pair types at an ITR. **D** Detail of the F2b and F4 read types. Note that the F2b is physical indistinguishable from an F2a read. An F4 read is read-pair where one read is complete telomere and the other is not. The complete end is comprised of TTAGGG

The Telomerecat length estimation method requires that all read pairs are sorted into four categories: F1, F2, F3 F4. Examples of each read type are given in Figure 9. Pseudocode for categorisation of reads is given in Supplementary Algorithm 3.

The read categorisation process is crucial to Telomerecat’s ability to filter interstitial reads. As we see in Figure 9, F2a are read pairs that straddle the boundary between telomere and the rest of the genome whereas F2b reads fall on one side of an ITR. We cannot directly observe the number of F2a or F2b read pairs; the orientation and sequence content of the read types are identical. However, we do know that, on average, within a sequencing experiment, there should be a corresponding F2b for each F4. Using this information we can deduce the amount of F2a reads.

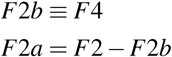

F4 reads give us an estimate of ITR reads, so subtracting F4 from F2 we are left with a count of reads F2 for which there was no corresponding F4. We posit that this is the count of reads on the boundary between telomere and subtelomere.

This method allows us to attain an estimate of F2a without filtering reads based on any upstream processing or any sequence structure beyond a distinction between “complete” and “incomplete” (see Supplementary Algorithm 3).

### Using cohort wide information to correct error in F2a counts

We observe that in some cases it is useful to normalise a cohort’s *F2a* count based on information from other samples in the batch. What follows is a method for adjusting F2a using a weighted average.

Let *C* be the total number of TELBAMs in a batch provided to Telomerecat. Such that subscript *c* represents a value relevant to any individual TELBAM. Let 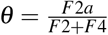 such that *θ^exp^* is the average *θ* observed across all TELBAMs in a cohort and 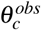 is the observed value of *θ* with in a particular TELBAM.

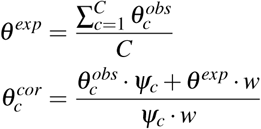

Where *w* is a predetermined weight of 3. *ψ* for any given TELBAM is obtained as follows.

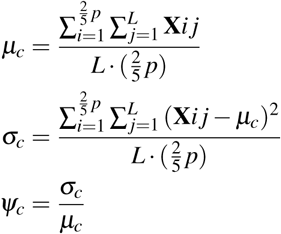

So it follows that the adjusted value of F2a is given as *θ^cor^* · (*F*2 + *F*4)

### Estimating length from read pair categories

The final step of the telomere length estimation process involves converting a ratio of *F*1: *F*2*a* read counts into an estimation of length. We achieve this by simulating telomere length under the observation of counts for F1, F2a and the fragment size. Psuedocode for the simulation is given in Algorithm 1

#### Algorithm 1 Telomerecat length estimation simulation algorithm

~~~
**function** LengthEstimation(F1,*F*2*a*)
         *τ* ← Arbitrary starting TL
         *μ, σ* ← Sample fragment mean and standard deviation
         **while** (*F*1′ ≠ *F*1)&(*F*2*a*′ ≠ *F*2*a*) **do**
           *F*1′,*F*2*a*′ ← *simulate*(*τ,F*1 + *F*2*a*,*μ,σ*)
           **if** *F*1′ < *F*1 **then**
              *τ* ← *τ* + *i*
           **else if** *F*1′ > *F*1 **then**
              *τ* ← *τ* − *i*
         **return** *τ*
~~~

### Batch effect correction when multiple sequencing platforms are used

Our observation has been that estimates from the HiSeqX platform are shorter on average than estimates from the HiSeq2000 platform. We have also observed that samples sequenced on the HiSeqX platform show lower scores in quality assessment. To account for this effect we propose that a mean correction should be applied to estimates from the HiSeqX platform.

## Acknowledgements

We thank Lawrence Bower for running bioinformatic pipelines, the Cambridge Cancer Research Fund and Hayley Whitaker for access to computing resources, and Zhao Ding for information regarding TelSeq. We also thank Chris Penkett for running bioinformatic pipelines and Hana Lango Allen and Ernest Turro for providing feedback on the technical replicates study.

This study makes use of data generated by the NIHR BioResource - Rare Disease. A full list of the investigators who contributed to the generation of the data is available from http://bioresource.nihr.ac.uk/rare-diseases/rare-diseases. Funding for NIHR BioResource - Rare Disease was provided for by the National Institute for Health Research.

We acknowledge Zhixin Qiu and colleges at Shanghai Institute of Biochemistry and Cell Biology for granting access to the HCC cell line data.

We acknowledge TwinsUK for providing WGS and mTRF telomere estimates. TwinsUK WGS data was generated by the UK10K Project. TwinsUK is funded by the Wellcome Trust, Medical Research Council, European Union, the National Institute for Health Research (NIHR)-funded BioResource, Clinical Research Facility and Biomedical Research Centre based at Guys and St Thomas NHS Foundation Trust in partnership with Kings College London.

JHRF, AGL and MLS were supported in this work by a Cancer Research UK Programme Grant to Simon Tavaré (C14303/A17197). Additionally, MLS was supported in this work by the European Community’s Seventh Framework Programme under grant agreement No. 305626 (Project RADIANT), and AGL by funding from the European Commission through the Horizon 2020 project SOUND (Grant Agreement no. 633974). We acknowledge the support of the University of Cambridge, Cancer Research UK and Hutchison Whampoa Limited

## Author contributions statement

J.F wrote and designed the algorithm, conducted the analysis and wrote the manuscript. M.S contributed to key elements of the algorithm. A.L conceived the concept for the algorithm and wrote the manuscript. All authors reviewed the manuscript.

